# Application of CRISPR-Cas12a technology in zebrafish to simulate ß-globin gene (HBB) mutations

**DOI:** 10.1101/2023.09.21.558926

**Authors:** Farheen Shafique, Shaukat Ali, Edward Balckburn, Mahreen ul Hassan

**Affiliations:** University of Azad Jammu and Kashmir; Government College University Lahore; The University of Sheffield; Shaheed Benazir Bhutto Women University

**Keywords:** CRISPR, Cas12a, MMEJ, Zebrafish, ß-globin, ß-thalassemia

## Abstract

**Background:** The human ß-globin (HBB) gene translated into the hemoglobin subunit beta (ß) of the hemoglobin protein. When mutated, it can lead to the blood disorder ß-thalassemia. Recently, the CRISPR-Cas12 technology has shown promising potential in treating different genetic abnormalities.

**Objectives:** This study aimed to evaluate the feasibility of zebrafish hbbe1.1 gene editing via CRISPR-Cas12a gene-technology.

**Methods:** In current study, thalassemic mutations were replicated in zebrafish to improve gene editing. the embryonic hemoglobin gene *hbbe1.1* was preferred over adult hemoglobin *hbß-a1* due to its early detection at larval stages. The CRISPR-Cas12 technology was utilised in combination with phosphorothioated “rescue template” (ssDNA L33P) to introduce base edits to the DNA sequence of zebrafish.

**Results:** These experiments indeed resulted in the alteration of a single amino acid at the protein level. With the help of this genetic editing method, we were able to generate a novel zebrafish strain that carried specific amino acid alterations resembling the pathogenic mutations found in HBB for ß-thalassemia disease. However, in several cases additional indels or base alterations were observed.

**Conclusion:** Our findings suggests that MMEJ double stranded break repair mechanism causes more knock-in events and germline inheritance than HR-mediated events. Modifying the technique could improve results by reducing MMEJ frequency.

## Introduction

ß-thalassemia is a common human blood disorder worldwide [1] characterised by hemolysis, iron overload, and inefficient erythropoiesis. The disease is caused by mutations in the ß-globin gene, leading to a malfunctioning ß-globin protein, which in turn leads to life-threatening anemia. Currently, frequent erythrocyte transfusions and chelation therapy are practised to reduce iron overload to alleviate symptoms [2].

The HBB gene encodes human ß-globin protein. which holds the haem molecule that binds oxygen and is therefore crucial for red blood cells, which transport oxygen throughout the human body [3]. The hemoglobin subunit ß adult 1 (*hbß-a1*) gene in zebrafish is similar to the *HBB* gene and performs the same function thus, zebrafish might be usable as a model for ß-thalassemia [4]. However, at embryonic stages, the human foetus express Gamma globin / *γ-globin* (HBG) gene, which is replaced by adult ß-hemoglobin (HBB) in later stages after birth [5]. Similarly, in zebrafish there is embryonic hemoglobin, encoded by the *hbbe* gene, which switches to the adult hemoglobin gene *hbß-a1*, 10-14 days after fertilisation [6].

Currently, the only curative treatment for ß-thalassemia is stem cell transplantation, which corrects the underlying genetic abnormality. Unfortunately, access to this treatment is limited by the capabilities of the local healthcare system, greater risks of graft-versus-host disease, death, transplant failure, and, most importantly, the lack of ideal donors [7]. With recent advances in the possibility of altering haematopoietic stem cell genes, gene editing has become a potential treatment for both hereditary and acquired disorders [8–9]. A robust and straightforward genome editing tool called Clustered Regularly Interspaced Short Palindromic Repeats (CRISPR) and CRISPR-associated protein 9 system (also known as CRISPR/Cas9) has recently been shown to open new avenues for exploring functional organisation of the genome and creating novel genetic variations [10]. Researchers can now directly modify or edit the function of DNA in organisms of their choice [11].

Since 2014, several labs have generated iPSCs from various types of cells recovered from individuals with ß-thalassemia and CRISPR-Cas9 technology was used to fix the mutated genes. In one study, reprogramming factors were used to generate iPSCs from skin cells from an individual who was heterozygous for the -28(A/G) mutation and the 4-bp (TCTC) deletion at codons 41 and 42 in exon 2. CRISPR-Cas9 paired with piggyback technology was used to repair these mutant stem cells [12]. It was shown that the HBB mutations in patient-derived iPSCs had been correctly repaired and that the cells retained their full pluripotency. Differentiated erythroblasts derived from these iPSCs exhibited normal HBB protein [13]. Another report showed effective repair of homozygous CD41/42 (-CTTT) mutations in iPSCs from a patient with -thalassemia using a combination of single-strand oligodeoxynucleotides (ssODNs) and CRISPR-Cas9 technology [12].

Despite many breakthroughs, CRISPR/Cas9 has downsides, mainly off-target activity, and unintended additional mutations at the on-target site [14–15]. Even the most efficient HDR-mediated CRISPR/Cas systems cannot avoid undesired mutations [16].

The accuracy of CRISPR/Cas-mediated genome editing may be increased by the recent discovery of additional programmable Cas proteins. For instance, Cas 12 extracted from Lachnospiraceae (LbCas12a) [17]. Cas12 has emerged as a Cas9-alternative for gene editing. The Cas12 protein uses crRNAs to cut single or double stranded DNA by utilizing its RuvC and NUC cleavage domains [18]. After PAM recognition, an R-loop forms. Cas12a cuts the non-target strand continuously, trimming it towards the PAM, and finally cut the target strand imprecisely. Due to the rate-limiting nature of R-loop creation for cleavage, Cas12a binding eventually results in target DNA cleavage. The Cas12 protein uses crRNAs to cut single or double stranded DNA by utilizing its RuvC and NUC cleavage domains [18]. After PAM recognition, an R-loop forms. The Cas12a cuts the nontarget strand continuously, trimming it towards the PAM, and finally cut the target strand imprecisely. Due to the rate-limiting nature of R-loop creation for cleavage, Cas12a binding eventually results in target DNA cleavage 19]. The Cas12 has some advantages over Cas9, including recognizing a more relaxed PAM sequence, having higher target specificity, targeting multiple sites within a single DNA molecule, and being smaller in size, which is advantageous for delivery into cells [20–23]. Both Cas12a and Cas9 as part of the CRISPR system are widely used for genome editing. However, in many cases, Cas12a performed better than Cas9 by making DSBs at different times to encourage HDR repair instead of both NHEJ and HDR [16].

The objective of this project was to investigate the feasibility of utilizing Cas12 in zebrafish to generate a specific mutation that induces an embryonic version of ß-thalassemia by targeting the zebrafish embryonic hemoglobin gene. The zebrafish was chosen because of its genetic similarity to humans and its small size to allow easier experimentation. The goal was to study the effects of these mutations on the organism once induced, and they might be exploited evaluate different gene editing techniques to correct them.

## Methods

### Zebrafish type and breeding

For the current project, a wild-type AB zebrafish strain (*Danio rerio*) wild-type strain AB or AB-LWT strain (London Wildtype) were used. The fish were kept on a 14-hour light cycle. Fertilised eggs were obtained by mating the fish described by Kimmel et al. (1995)[24]. Embryos were raised in 1X E3 embryo medium prepared by diluting 16.5 ml 60X stock in 1 L double deionized water after which 100 μl of methylene blue was added [25]. Embryos were kept at a maximum temperature of 28.5°C maximally 50 in a 10 cm Petri dish upto 5dpf unless otherwise indicated. The staging was done by following by Kimmel et al. (1995)[24].

### Single guide RNA (sgRNA) design

A 20bp long targeting guide was selected using CHOPCHOP v2 [26]. Targets starting with ’GG’ were preferred using a T7 promoter. The PAM sequence (NGG) adjacent to the CRISPR site was also annotated, to ensure that it would be excluded from the guide sgRNA design. To facilitate the identification of mutants through restriction digestion, preference was given to CRISPR target sites that overlapped with enzyme restriction sites, as determined by the NEB cutter software.

### System Design to Generate Transgenic Zebrafish hbbe1.1 (L33P)

As described by Renaud et al. (2016) and Richardson et al. (2016), the ssDNA repair template containing a mutated PAM site (S1 Table) to avoid recurring nuclease activity was constructed for effective HDR-mediated repair at the target location and ordered from Interactive DNA Technologies (IDT) [27–29].

A 20 bp gRNA was designed with complementarity to the non-target strand as shown in Table 1. A 3’-uridylated (U8) tail was added to enhance INDEL efficiency as suggested by Bin Moon et al., (2018) [30]. The generic Cas12a binding region was designed and ordered from Sigma-Aldrich^®^.

### CRISPR Microinjection method for gene delivery

To prepare the CRISPR-Cas12a microinjection solution, 1 μL CrRNA (200μM) (Sigma), 0.5 μL Cas12a (100 μM) (NEB) and 0.2 μl of 10X NEB 2.1 buffer were mixed gently in a PCR tube and incubated at 25°C for 15 minutes in a Thermal Cycler (Bio-Rad T100) and then 1 μL ssDNA repair template [IDT] and Phenol red (0.3 μL) were added to the mixture. MilliQ was added to 4 ul and stored on ice until use. As a negative control, 1 μL of ssDNA repair template (100 μM) was mixed with 3 μL of MilliQ. 0.5nl of the mixtures were directly injected into the yolk of freshly layed zygotes using a glass capillary needle [TW120-4) (Kwik-Fil, World Precision Instruments, Inc., Hertfordshire, UK)] with the help of a pedal controlled microinjector (PV820 Pneumatic PicoPump, WPI).

### Mutation-specific Genotyping through touch down PCR

DNA was extracted from embryos using the HOTSHOT extraction method (S1 Protocol) while larval DNA was extracted using the Proteinase K extraction method (S2 Protocol). After 24h genomic DNA was analyzed through gene amplification using a forward and a mutation specific reverse primer by standard PCR (Bio-Rad) using Firepol Taq Polymerase [BioDyne]. Visualization of PCR products after agarose gel electrophoresis was performed using a Benchtop UV Transilluminator [UVP]. Mutation specific primers were used to amplify and screened potential positive fish unclear, and Sanger sequencing [Azenta Life Sciences].

### Genome Analysis and Manipulation

Genomic sequence data for both humans (GRCh38.p13) and zebrafish (GRCz11) was retrieved from ENSEMBL genome browser. SnapGene® software 5.2.5 (from Dotmatics; available at snapgene.com) was used for in silico sequence analysis and modification (Snapgene 5.2.1). CHOPCHOP v2 web tool was used to assess the quality of guide sequences [26]. Primers were designed using Primer3 and validated with NCBI Primer-BLAST to guarantee specificity [31–32].

### Generation of mutant lines by CRISPR genome editing

After confirmation that a CRISPR guide could create mutations, at least 50 injected G0 embryos were raised to adulthood. Adult G0 fish were outcrossed with ABWT fish, and their progeny was screened for germline transmission by PCR and restriction analysis followed by Sanger sequencing.

### Bright-field imaging

Bright-field images were taken via the Lecia M165 FC microscope using LAS V4.9. The imaging was performed by scanning a 1024 x 1024 pixel resolution. The zebrafish were treated with tricaine, and 3% Methylcellulose was used to mount the zebrafish in the correct side-view orientation.

## Results

The project had the primary objective of developing and optimising gene editing techniques. To achieve this, the embryonic hemoglobin gene was chosen as the target gene for mutation. Specifically, the objective was to replicate thalassemic mutations within the model organism of zebrafish. This was a deliberate choice, as the zebrafish has emerged as an increasingly popular model organism for studying human diseases due to their high genetic similarity to humans and their small size, which facilitates easier experimentation. By replicating these thalassemic mutations in zebrafish, it would be possible to study the effects of these mutations on the organism, as well as test the efficacy of various gene editing techniques to correct these mutations. Ultimately, the findings of this project could have significant implications for the development of gene editing therapies for human genetic diseases.

### Characterisation of the hbbe1.1 gene in zebrafish

The zebrafish embryonic hemoglobin *hbbe1.1* was preferred on adult hemoglobin *hbba* gene due to rapid growth of fish embryos and early signs of disease (loss of blood colour due to malfunctioning hemoglobin) in fish larvae i.e., starting from day 3 post fertilization. This time window would make it possible to induce and analyse ß-globin mutations without causing secondary defects due to oxygen insufficiency.

The *hbbe1.1* gene in zebrafish was recovered by ENSEMBL searches. The gene was also scanned through NCBI to ensure that there were no additional orthologues. The alignment of the sequence of the human protein (NP_000509.1) and zebrafish (NP_932339.1) protein indicated a share of 51.72% sequence similarity (S1 Fig).

### Target site selection in the hbbe1.1 gene

the zebrafish *hbbe1.1* gene (S2 Fig) was targeted for CRISPR/Cas12a mutagenesis. A homology-directed repair (HDR) approach was used to produce mutant variants [*Hbbe1.1 c.332 CTG(Leu)>CCG(Pro)(p.Leu33Pro*)] similar to the human HBB variant [*HBB:c.92 G>C or IVS-I (−1) AG^GTTGGT->AC^GTTGGT (p.Arg30Thr*)] causing ß-thalassemia reported by Yasmeen et al. (2016)[33].

### CRISPR selection

In the current study we use direct delivery of Cas12a protein for efficient CRISPR mediated gene editing. Injecting an active Cas12a protein along with gRNA makes a very efficient RNP complex for targeted gene modification with minimum off-target effects.

For the development of individual CRISPR systems it was important to gather detailed information about the genomic sequence and main protein structure of the *hbbe1.1* variation (L33P) by utilising the ENSEMBL genome browser [34]. The present investigation employed the ENSEMBL database to elucidate the orthologous relationship between the hbbe1.1 gene and its human homolog. Subsequently, the ENSEMBL ID was utilized to predict potential cleavage sites for *hbbe1.1* (Cas12a) through the utilization of the bioinformatics tool CHOPCHOP, which is commonly used to design guide RNAs. The purpose of this analysis was to identify candidate disease-associated variable codons that lie in close proximity to a CRISPR target location, with a maximum distance of 10 base pairs [35].

### System design for CRISPR-Cas12a mediated base edits (*hbbe1.1*) at codon 33, CTG (leucine) to CCA (proline) ATC

The CRISPR-Cas12a ssDNA repair template requires a PAM-distal arm containing the *hbbe1.1* target site *hbbe1.1* (codon 33 in this case) and some additional silent mutations downstream to increase mutation-specific binding. As described by Miao et al. (2019), mutations should not exceed <10 bp from the Cpf1 cleavage site. To maintain the stability of the rescue template and to avoid DNA degradation by exonucleases, the last two bases are phosphorylated at the 5 and 3 ends of ssDNA [35, 27]. The PAM site of the rescue template must be mutated to avoid recurrent nuclease activity [29].

The gRNA (Sigma-Aldrich) design (Table S9) requires a 20 bp complementary sequence to the non-target strand of the gene of interest; A 3’-uridylated (U8-) tail for increased INDEL efficiency (Bin Moon et al. 2018)^30^, and a generic region (5’-UAGGUAAUUUCUACUAAGUGUAGAU–N20–UUUUUUUUU-3’) for the Cas12a protein (S2 Fig). The target site was aligned with the human adult hemoglobin HBB gene with a similar mutation CTG>CGG (Leucine to Arginine) at codon 91 (Fig 1) reported by Ali et al., (1988) in Canadian ethnicity [36].

**Fig 1.**
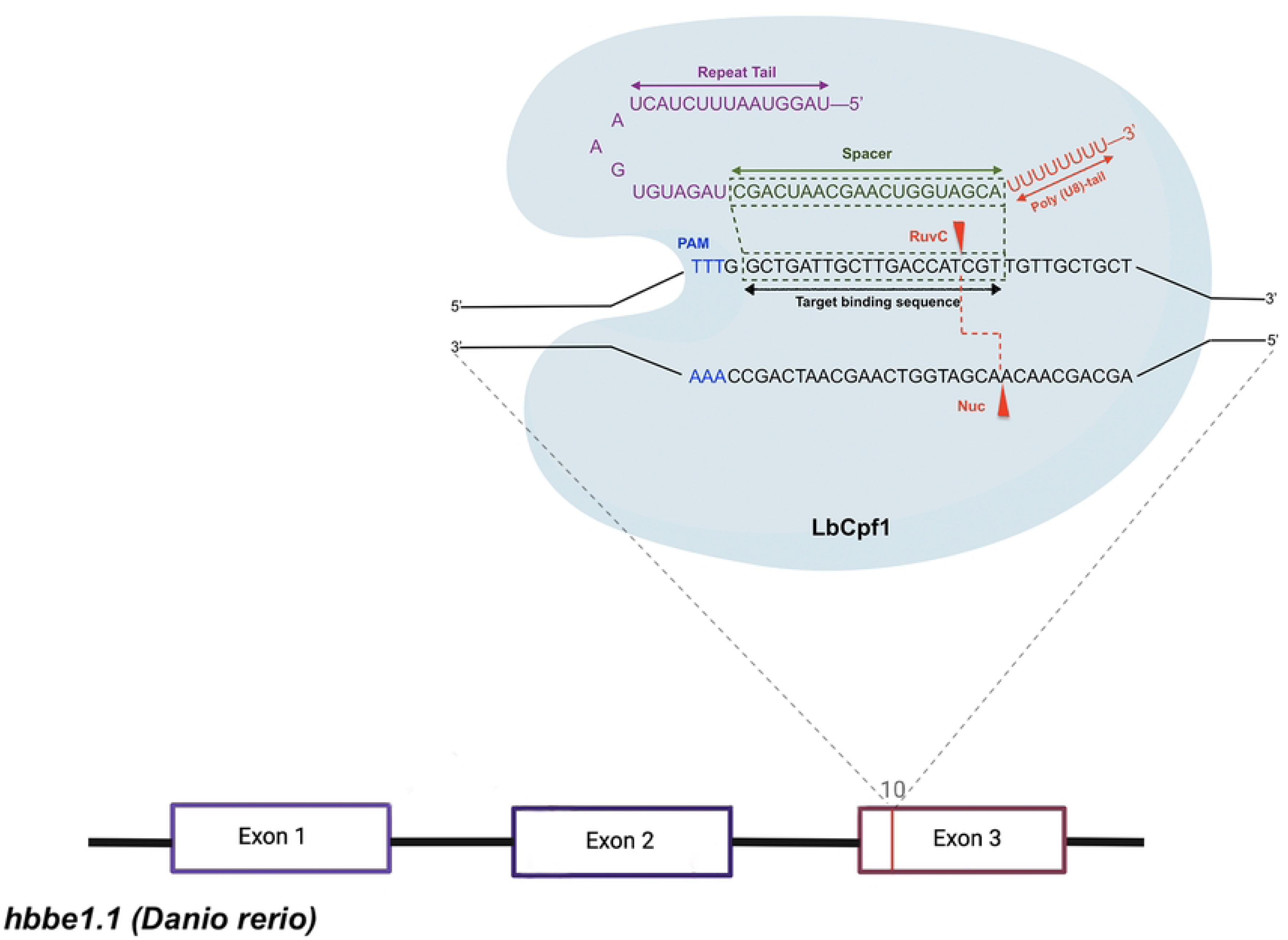
Schematic representation of the *hbbe1.1* gene for mutation induction at codon 33. The Nuc and RuvC domains (red) break DNA to form a sticky end.

The gRNA to target the hbbe1.1 codon 33 for base substitution was designed through an online CRISPR gRNA design tool CHOPCHOP for gene repression through Cas12a nuclease. The target was selected with a 50% GC content and only one off-target (A→G) at position chr3: 5123086 (S3 Fig).

The repair template *hbbe1.1-L33P* to induce the L33P mutation in hbbe1.1 at codon 33 inside the zebrafish genome and the gRNA for Cpf1-L33P are given in Table S1.

PCR primers were designed using Primer3 online tool to monitor L33P gRNA and validate the hbbe1.1-L33P rescue template integrated into the zebrafish genome. Mutation-specific PCR primers were also designed to differentiate between mutant and wild-type sequences. Additionally, a set of sequence primers was designed for sequence analysis (see S2 Table).

### Integration of *hbbe1.1*-L33P in G0 zebrafish

Microinjection was prepared to inject 1nl of a mixture of Cas12a, rescue template L33P, gRNA and phenol red in freshly laid zebrafish embryos. To check the efficiency of guide some of the injected embryos were kept for DNA extraction and gene amplification. The guide efficiency primers were used to amplify the DNA (S2 Table).

For the validation of whether the L33P repair template is incorporated into the zebrafish genome, special primers were designed that flank the target area. On day 3, at about 120 hpf, some larvae were sacrificed to extract DNA for PCR. A set of validation primers (S2 Table) was designed and used to amplify the targeted DNA.

The PCR results showed the expected binding of the validation primers in the 196 bp position on the agarose gel (S4 Fig). The forward validation primer flanked the mutated sequence, while the reverse primer overlapped the 5’ end of the rescue sequence. No amplification was found in the water control and wildtype un injected embryonic DNA wells as water control lacked the targeted sequence and the wildtype embryos were not injected at all. This suggests that the *hbbe1.1*-L33P validation primers *hbbe1.1*-L33P were able to bind to the mutated sequence only. Another higher band could also be seen above the 196bp band which suggests addition of bases at cut site which might be due to the MMHJ repair mechanism acquired by the cells after DBS. The heteroduplex formation might be simply because of variation in temperature or nonspecific binding of primers to the DNA template, generating a mixture of PCR products. On day 4 the zebrafish larvae were again examined under the microscope. A reduction in redness of the blood in heart region was observed confirming that the guides were active (Fig 2). A batch of about 80 wildtype embryos were kept from the same parents as a control group, to control for abnormalities unrelated to the injection. After confirmation of the incorporation of the mutation at the right site through PCR, mutant embryos with no known abnormalities other than colourless blood were raised.

**Fig 2.**
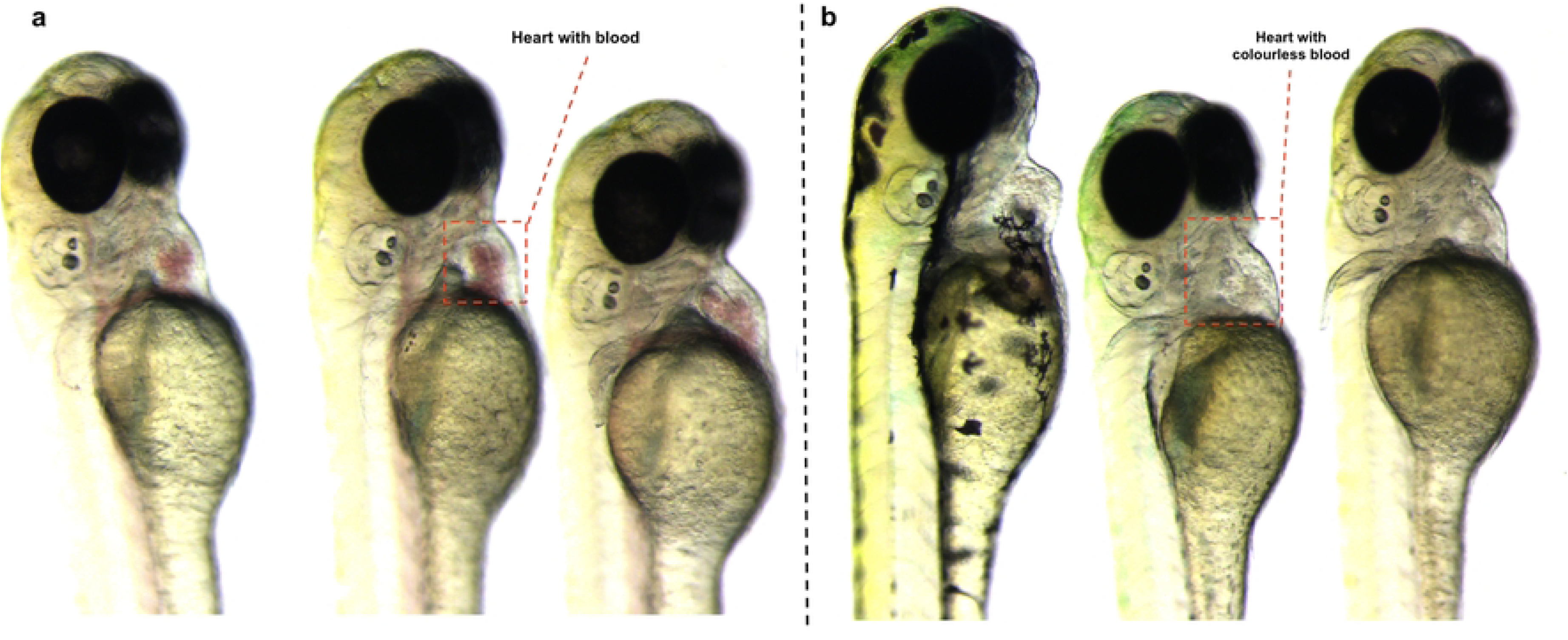
Comparison of blood colour in the heart region of zebrafish larvae. (a) healthy wildtype zebrafish larvae. **(b)** mutant zebrafish larvae injected with hbbe1.1 - Cas12a. The heart region contains colourless blood due to malfunctioning hemoglobin.

### Generation of hbbe1.1 L33P F1 transgenic zebrafish

To study the germline transmission of the knocked in *hbbe1.1-L33P* rescue template, larvae injected together with gRNA and Cas12a protein were sent to the aquarium of the University of Sheffield, UK, for raising. The larvae were kept in a techniplast aquarium system and fed with artemia and rotifers regularly for 12 weeks until sexual maturity. These injected G0 fish might be mosaic for any induced mutations, therefore they needed to be outcrossed with wildtype fish (Fig 3) to observe if the G0 founders were able to transfer mutations of interest to the next generation or not. After 3 months the healthy-looking adult male and female fish were selected and mated with wildtype zebrafish in mating boxes as described by Westerfield (1993) [37]. Around 12 pairs of fish were mated among which 8 pairs laid eggs. The embryos were carefully collected and incubated at 28 °C. After a few hours, embryos were examined under the microscope and sorted to remove unfertilised and abnormally developing eggs. A few embryos were selected for DNA extraction.

**Fig 3.**
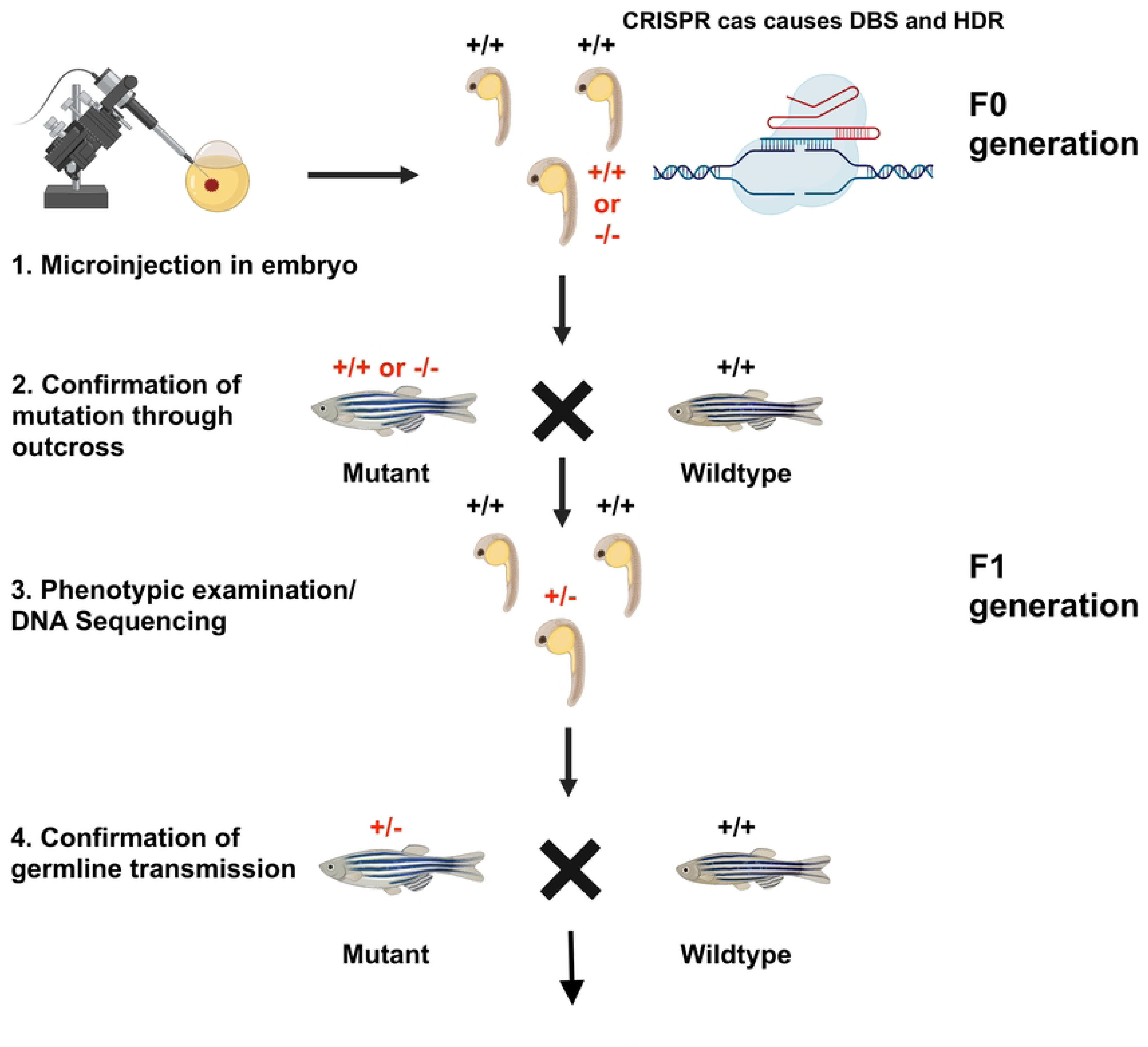
Schematic representation to describe generation of hbbe1.1 L33P transgenic zebrafish

### PCR assay for hbbe1.1 L33P mutation screening

After DNA extraction a PCR assay was performed to screen for the L33P mutation in the embryos from the G0 parents along with a wildtype genomic DNA and water as negative controls. First, the validation primers for *hbbe1.1 L33P* (S2 Table) were used to amplify the *hbbe1.1* gene that carries the L33P mutation. The same primers were used to verify HDR-mediated mutation within the zebrafish genome of the G0 generation after injections (S4a Fig). The PCR results confirmed the presence of the *hbbe1.1*-L33P mutation (196bp band) in some of the embryos as shown in S5 Fig. A double band was also observed in the DNA of the embryo of pair 3 which suggested an additional insertion event had occurred, which could be due to the alt-NHEJ event at the cleavage site.

The mutation was also confirmed through guide efficiency primers (S6a Fig). The primers would amplify a product of 132 bp. The samples 2 and 4 showed a band around 150 bp while the sample containing water control showed no band at all (S6b Fig). A smearing of band could be seen in most of the samples especially in samples 8, 9 10 and 11 which is the sign of mosaicism and suggests that there might be an INDEL event that occurred during Cas12a nuclease activity, which led to MMHJ by the cell to repair the DBS.

Before sending mutant F1 embryos for raising, it was necessary to screen them for the *hbbe1.1-* L33P mutation in the targeted region. For this reason, a set of mutation-specific primers (S2 Table) was designed to screen embryos for the targeted mutation. The mutation specific primers were designed in such a way so that they could only bind to the mutation (L33P) carrying gene inside the zebrafish genome (S7a Fig). A PCR would amplify a band of 139 bp if the mutation had been integrated at the target position (S7b Fig).

### Detection of CRISPR-Cas12a mediated mutation in the *hbbe1.1*

#### gene sequence

The PCR positive transgenic and wildtype zebrafish embryos were amplified by using PCR with *hbbe1.1*-L33P specific primers to detect the mutation. PCR amplification was carried out with specific primers that flank the sgRNA target site in the hbbe1.1 gene. A 139 bp fragment appeared corresponding to the amplified *hbbe1.1*-L33P gene in zebrafish and separated in 3% agarose gel electrophoresis (S6b Fig). The PCR amplified products were purified by the Exo-sap method and sent to GENEWIZ, Azenta Life Sciences, Hertfordshire, United Kingdom, for Sanger sequencing. DNA sequencing was performed with a specially designed L33P sequence reverse primer (S2 Table). Sequencing was carried out on twenty potential *hbbe1.1*-L33P mutation carrying transgenic zebrafish embryos from 2 different G0 parents (10 from each pair) with same mutation (Fig 4).

**Fig 4.**
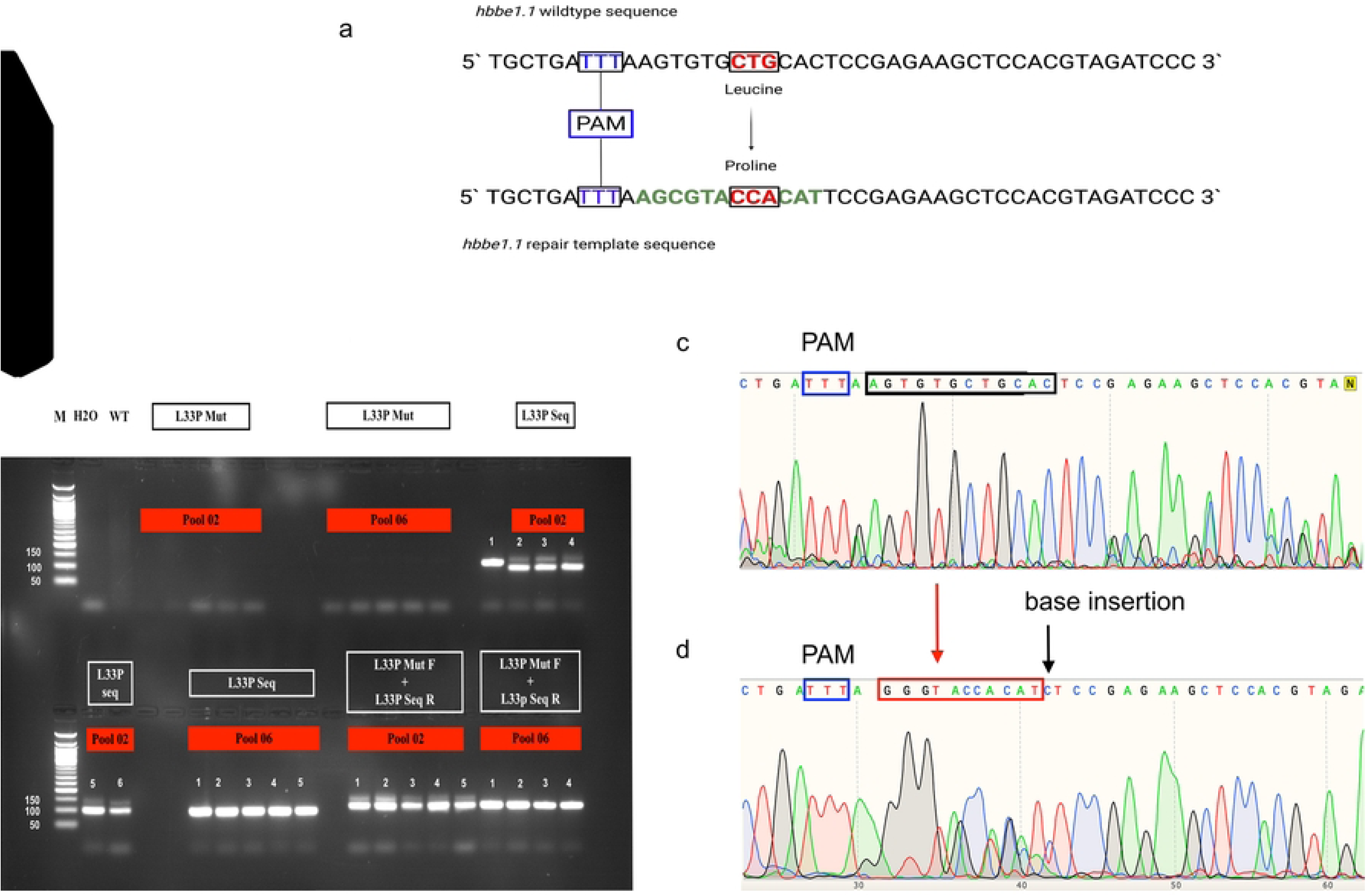
Sanger sequencing results of G1 *hbbe1.1-L33P* transgenic zebrafish. (a) Wild-type genomic sequence of the hbbe1.1gene and the hbbe1.1-L33P repair template. (b) The gel image showing potential samples found positive after amplification with different sets of mutation-specific and sequence primers listed in Table 3.15. The pool names in red represent the progeny from pair 2 and 6. (c) DNA sequence chromatogram showing the wild-type sequence of the hbbe1.1 gene. (d) DNA sequence chromatogram of hbbe1.1-L33P gene integrated in the G1 transgenic zebrafish genome from pool 2 and 6.

The sequence results revealed that the intended mutation CTG→CCA was inserted at the right position in the zebrafish genome but with some base substitutions (AGC→GGG) at the targeted mutation site. Furthermore, a base insertion ‘C’ was observed in the samples just next to the CCA mutation. The sequence results clearly indicated that cells of G0 fish acquired a type of MMHG repair mechanism during Cas12a nuclease activity. The same base substitution was found in all samples from pair 6. Although only two samples from pair 2 had shown the in-frame knock-in at the targeted site. The experiment was repeated with all the potential mutation-carrying fish.

After carefully screening all the *hbbe1.1*-L33P founder fish by out-crossing them with wildtype zebrafish it was observed that the 4 out of 10 (40 %) fish from pool 6 and 2 out of 10 (20 %) fish pool 2 showed in-frame knock-ins inherited from G0 parents at the targeted position inside their genome.

## Discussion

There have been numerous studies on gene editing techniques for the treatment of thalassemia. However, the project in question aimed to develop and optimize gene editing techniques using the zebrafish model organism. Here we tried to develop and optimize gene editing techniques by replicating thalassemic mutations within the model organism of zebrafish. The embryonic hemoglobin gene *hbbe1.1* was chosen as the target gene for mutation due to the rapid growth of fish embryos and early signs of disease in fish larvae. The objective was to induce and analyze ß-globin mutations without causing secondary defects due to oxygen insufficiency. The study used direct delivery of Cas12a protein for efficient CRISPR-mediated gene editing with minimum off-target effects. The study utilized an HDR approach to produce mutant variants [*hbbe1.1* c.332 CTG(Leu)>CCG(Pro)(p.Leu33Pro)], similar to the human HBB variant causing β-thalassemia. The findings of this project have significant implications for the development of gene editing therapies for human genetic diseases.

While there is limited information available on studies specifically targeting the *hbbe1.1* gene in zebrafish, several studies have focused on the use of CRISPR/Cas systems in zebrafish to model human genetic diseases. For example, a study by Morales and Wingert (2017) used the CRISPR/Cas9 system to create a zebrafish model of polycystic kidney disease, while another study by Tessadori et al. (2018) used the CRISPR/Cas9 system to generate a zebrafish model of Cantú syndrome (CS) [38–39]. Additionally, several studies have used zebrafish models to study anemia, such as the study by Bowman et al. (2017) that used sf3b1 gene mutations responsible for macrocytic anemia in zebrafish [40]. A study by Traxler et al. (2016) used CRISPR-Cas9 to induce mutations in human hematopoietic stem cells to correct the β-globin gene responsible for sickle cell disease and β-thalassemia [41]. The study demonstrated the feasibility of correcting disease-causing mutations in human hematopoietic stem cells using CRISPR-Cas9, which has implications for the development of gene editing therapies for human genetic diseases. Another study by Dever et al. (2016) used CRISPR-Cas9 to correct the β-thalassemia mutation in human CD34+ cells. The study demonstrated that the CRISPR-Cas9 system can specifically correct the β-thalassemia mutation, resulting in increased β-globin expression and reduced γ-globin expression [42].

The molecular analysis of the modified *hbbe1.1* gene integrated in the zebrafish suggested that the strategy for base substitution employed in current study could be improved by chemically modifying the repair template to inhibit alt-NHEJ and shift toward HDR-mediated repair as described by Aksoy et al. (2019) [43]. The findings of this project may help in the development of gene editing therapies for hereditary disorders.

## Conclusion

It was concluded from these results that the knock-in events mediated by MMEJ double stranded break (DSB) repair mechanism seems to be more prevalent and result in germline inheritance than HR-mediated events. This study showed that direct delivery of Cas12a protein for efficient CRISPR-mediated gene editing with minimal off-target effects is possible. A slight modification in this technique could reduce the frequency of MMEJ events that would lead to significantly improved results.

## Acknowledgements

We thank Dr. Fredericus Van Eeden for supervising the project and providing lab facilities. We are also thankful to the Aquarium staff members of the Biological Services Aquarium at Bateson centre who cared for our zebrafish at the University of Sheffield, United Kingdom.

## Author Contributions

FS was responsible for designing the study, writing the first draft, conducting the research, extracting, and analysing data, and interpreting results. SA was responsible for the review of study and supervision. EB helped in analysing data and interpreting results. MUH helped in editing of final draft.

## Ethical Approval

The procedures were regulated by following the UK Home Office Animals (Scientific Procedures) Act 1986 guidelines under project licence PC39B259E (Dr. Fredericus Van Eeden).

## Data Availability

Data could be provided upon request.

